# FoxO3a and miR-34a-3p Are Involved in Oxidative Stress-Induced Dysfunction of Human Endothelial Progenitor Cells

**DOI:** 10.64898/2026.07.03.736301

**Authors:** Ziyu Lin, Jifang Ban, Yuqiang Wang

## Abstract

**Background:** Endothelial progenitor cells (EPCs) contribute to endothelial repair and neovascularization, and EPC dysfunction is closely associated with oxidative stress-related vascular injury. Forkhead box O3a (FoxO3a) regulates cellular stress responses, whereas miR-34a has been implicated in endothelial dysfunction, senescence, and apoptosis. However, the relationship between FoxO3a and miR-34a-3p in oxidatively injured EPCs remains incompletely defined.

**Objective:** This study investigated the role of FoxO3a in H_2_O_2_-induced EPC dysfunction and examined whether miR-34a-3p directly interacts with the FoxO3a 3′ untranslated region (3′UTR).

**Methods:** Human umbilical cord blood-derived EPCs were identified by DiI-ac-LDL uptake, FITC-UEA-1 binding, and the expression of EPC-related markers. Oxidative stress was induced by H_2_O_2_. Cell viability, apoptosis, and angiogenic capacity were evaluated using CCK-8 assay, Annexin V/7-AAD flow cytometry, and Matrigel tube formation assay, respectively. FoxO3a expression was modulated using adenoviral overexpression or knockdown vectors, and miR-34a was modulated using mimics or antagomir. FoxO3a and miR-34a expression levels were detected by Western blot and qPCR. A dual-luciferase reporter assay was used to verify the interaction between hsa-miR-34a-3p and the FoxO3a 3′UTR.

**Results:** H_2_O_2_ reduced EPC viability, increased apoptosis, and impaired tube formation in a concentration-dependent manner. H_2_O_2_ increased FoxO3a protein abundance and miR-34a expression, whereas FoxO3a mRNA did not change markedly. FoxO3a overexpression aggravated, whereas FoxO3a knockdown partially alleviated, H_2_O_2_-induced EPC dysfunction. Similarly, miR-34a mimics further suppressed EPC viability and tube formation, while miR-34a antagomir exerted a protective effect. Dual-luciferase reporter analysis showed that hsa-miR-34a-3p significantly reduced the activity of the wild-type FoxO3a 3′UTR reporter, while mutation of the predicted binding site abolished this suppression.

**Conclusion:** FoxO3a and miR-34a participate in oxidative stress-induced EPC dysfunction. The dual-luciferase data demonstrate that hsa-miR-34a-3p directly targets the FoxO3a 3′UTR, suggesting the presence of miR-34a-3p-mediated post-transcriptional feedback within the FoxO3a-related stress-response network in EPCs.

## 1. Introduction

Endothelial progenitor cells (EPCs) are circulating progenitor cells with endothelial differentiation potential and paracrine activity. They contribute to endothelial repair, postnatal neovascularization, and vascular homeostasis. Under pathological conditions, EPCs can migrate to sites of endothelial injury and participate in repair through endothelial differentiation and the release of pro-angiogenic and anti-inflammatory mediators. EPC dysfunction is commonly observed in cardiovascular and metabolic diseases, including atherosclerosis, hypertension, diabetes, and ischemic vascular disease, and is typically characterized by reduced viability, increased apoptosis, impaired migration, and attenuated angiogenic capacity [1,2].

Oxidative stress is one of the major mechanisms underlying EPC dysfunction. Excessive reactive oxygen species (ROS) disrupt cellular redox homeostasis, damage proteins, lipids, and nucleic acids, and activate apoptosis-related signaling pathways. In vascular tissues, ROS accumulation is closely associated with endothelial dysfunction, reduced nitric oxide bioavailability, inflammatory activation, and abnormal vascular remodeling [3]. Hydrogen peroxide (H_2_O_2_) is widely used to establish in vitro oxidative stress models because it penetrates cell membranes and induces intracellular oxidative injury. Previous studies have shown that H_2_O_2_ exposure reduces EPC survival and promotes apoptosis, indicating that oxidative stress directly compromises EPC viability and function [4].

Forkhead box O3a (FoxO3a), a member of the FoxO transcription factor family, participates in stress responses, metabolism, inflammation, autophagy, cell-cycle control, antioxidant defense, and apoptosis [5]. FoxO3a has context-dependent biological effects. Under mild or transient stress, FoxO3a may support adaptive antioxidant responses and cellular homeostasis; however, under sustained or severe stress, excessive FoxO3a activity can promote apoptosis through pro-apoptotic mediators such as BIM. In EPCs, H_2_O_2_-induced impairment of viability has been reported to involve a FoxO3a-dependent mechanism [4]. SIRT1 has also been shown to protect EPCs from oxidative stress-induced apoptosis by modulating FoxO3a protein stability and activity [6]. These findings indicate that FoxO3a may serve as an important molecular link between oxidative stress and EPC injury.

MicroRNAs are small non-coding RNAs that regulate gene expression predominantly at the post-transcriptional level and participate in cell proliferation, apoptosis, senescence, inflammation, angiogenesis, and vascular remodeling [8]. miR-34a is a stress-responsive microRNA that has been implicated in endothelial senescence, apoptosis, and impaired angiogenesis [7,15]. Because these processes are closely related to EPC-mediated vascular repair, abnormal miR-34a expression may have important consequences for EPC function under oxidative stress.

Sirtuin 1 (SIRT1) is an NAD^+^-dependent deacetylase with protective roles in vascular cells. It regulates redox homeostasis, mitochondrial function, inflammation, apoptosis, and angiogenesis [19,20]. SIRT1 signaling has been reported to improve EPC function under pathological conditions [9], and miR-34a is widely recognized as an upstream post-transcriptional regulator of SIRT1 in several vascular contexts [7,15]. Therefore, miR-34a-related signaling may contribute to EPC injury by converging on SIRT1-associated protective pathways. However, direct mechanistic conclusions regarding SIRT1 require specific rescue experiments and should be distinguished from evidence demonstrating miR-34a/FoxO3a interaction.

Previous evidence also suggests that FoxO3a and miR-34a may be functionally connected. Natarajan et al. reported that FoxO3 increased miR-34a expression in palmitate-induced cholangiocyte lipoapoptosis [10]. Conversely, miRNAs frequently regulate FoxO family members through 3′UTR binding. Whether hsa-miR-34a-3p directly interacts with the FoxO3a 3′UTR in the context of oxidative stress-related EPC dysfunction remains unclear.

In this study, we established an H_2_O_2_-induced EPC injury model and evaluated EPC viability, apoptosis, and tube formation. We further examined the effects of FoxO3a overexpression or knockdown and miR-34a mimic or inhibitor transfection on EPC function. Finally, a dual-luciferase reporter assay was performed to verify the direct interaction between hsa-miR-34a-3p and the FoxO3a 3′UTR. The study aimed to clarify the roles of FoxO3a and miR-34a in oxidative stress-induced EPC dysfunction and to provide experimental evidence for a FoxO3a/miR-34a regulatory network in EPC injury.

## 2. Materials and Methods

### 2.1 Isolation and culture of EPCs

Human umbilical cord blood was obtained from four independent donors after informed consent and approval by the institutional ethics committee. Mononuclear cells were isolated by density gradient centrifugation and washed twice with phosphate-buffered saline (PBS, pH 6.8). Cells were resuspended in EGM-2MV medium (Lonza, USA) supplemented with 5% fetal bovine serum and seeded in culture plates.

After 3 days, non-adherent cells were removed, and adherent cells were maintained in fresh medium. The medium was replaced every 3 days. EPC colonies were passaged when necessary, and cells at passages 3-4 were used for subsequent experiments.

### 2.2 Identification of EPCs

Cultured EPCs were identified by DiI-ac-LDL uptake, FITC-UEA-1 binding, and immunofluorescence staining of EPC-related surface markers. Briefly, EPCs were incubated with DiI-ac-LDL at 37°C for 4 h, washed with PBS, and then incubated with FITC-UEA-1 at 37°C for 1 h. After washing, nuclei were counterstained with DAPI. Fluorescence images were captured using an inverted fluorescence microscope.

For surface-marker staining, EPCs were fixed with 4% paraformaldehyde and blocked with normal serum or blocking buffer. Cells were incubated with primary antibodies against CD34, CD133, and VEGFR-2 overnight at 4°C, followed by fluorescence-conjugated secondary antibodies. Nuclei were counterstained with DAPI and observed under a fluorescence microscope.

### 2.3 Adenoviral construction and transduction

Recombinant adenoviruses expressing FoxO3a (OE-FoxO3a), FoxO3a shRNA (sh-FoxO3a), and corresponding negative controls (OE-NC and sh-NC), each tagged with GFP, were obtained from Han Heng (China).

EPCs were transduced with adenoviruses at multiplicities of infection (MOIs) ranging from 5 to 120 for 2 h, followed by replacement with fresh complete medium. Cells were incubated for an additional 48 h before analysis. Transduction efficiency was evaluated by GFP fluorescence using an inverted fluorescence microscope (Leica DMI4000B). Based on the balance between transduction efficiency and cytotoxicity, an MOI of 80 was selected for sh-FoxO3a, and an MOI of 100 was selected for OE-FoxO3a and the corresponding controls.

### 2.4 miR-34a mimic and inhibitor transfection

miR-34a mimics, miR-34a inhibitors (antagomir), and their corresponding negative controls were purchased from GenePharma (Shanghai, China). EPCs in the logarithmic growth phase were transfected using EndoFectin IMAX (GeneCopoeia, USA) according to the manufacturer’s instructions. Transfection efficiency was assessed by qPCR.

### 2.5 Cell viability assay

Cell viability was assessed using the CCK-8 assay (APE BIO). EPCs were seeded in 96-well plates in triplicate and treated with different concentrations of H_2_O_2_ for 24 h. Optical density was measured at 490 nm using a microplate reader (Thermo Fisher Scientific).

### 2.6 Apoptosis analysis

Apoptosis was evaluated using an Annexin V-APC/7-AAD apoptosis detection kit (Keygentec). After treatment, EPCs were collected, washed with PBS, and stained according to the manufacturer’s instructions. Samples were analyzed by flow cytometry, and apoptosis rates were quantified using Multicycle software (Beckman Coulter).

### 2.7 Western blot analysis

Total protein was extracted using RIPA buffer supplemented with protease inhibitors. Protein samples were separated by SDS-PAGE and transferred onto nitrocellulose membranes. Membranes were incubated with primary antibodies against FoxO3a (12829S; Cell Signaling Technology, USA) and β-actin (3700S; Cell Signaling Technology, USA), followed by HRP-conjugated secondary antibodies. Signals were detected using ECL reagent (Millipore).

### 2.8 Quantitative real-time PCR

Total RNA was extracted using TRIzol reagent (Thermo Fisher Scientific). cDNA synthesis was performed using an Evo M-MLV RT Kit. Quantitative PCR was conducted using SYBR Green Master Mix on a 7500 Real-Time PCR system. Relative expression levels were calculated using the 2−ΔΔCt method and normalized to β-actin or U6, as appropriate.

### 2.9 Tube formation assay

Matrigel (80 μL per well) was added to 24-well plates and incubated at 37°C for 30 min to allow polymerization. EPCs were seeded onto the Matrigel and incubated for 24 h. Capillary-like structures were observed under a microscope, and the number of closed networks was quantified in four randomly selected fields.

### 2.10 Dual-luciferase reporter assay

The FoxO3a 3′UTR fragment containing the predicted hsa-miR-34a-3p binding site was cloned into the pmirGLO luciferase reporter vector to generate the wild-type construct (pmirGLO-FoxO3a-3′UTR-WT). A mutant construct (pmirGLO-FoxO3a-3′UTR-MUT) carrying disrupted seed-matching nucleotides was generated using site-directed mutagenesis. The constructs were verified by restriction enzyme digestion and sequencing.

293T cells were seeded in 96-well plates and co-transfected with WT or MUT reporter plasmids together with hsa-miR-34a-3p mimic or negative control using Lipo3000 reagent. Where indicated, cells were exposed to H_2_O_2_ after transfection. At 48 h after transfection, luciferase activity was measured using the Promega Dual-Luciferase Reporter Assay System. Firefly luciferase activity was normalized to Renilla luciferase activity.

### 2.11 Statistical analysis

All experiments were performed using at least three independent biological replicates unless otherwise indicated. Data are presented as mean ± standard deviation (SD). Normality was assessed using the Shapiro-Wilk test, and homogeneity of variance was evaluated using Levene’s test. Differences between two groups were analyzed using unpaired Student’s t-test. For comparisons among multiple groups, one-way or two-way ANOVA followed by Tukey’s post hoc test was applied. A value of P < 0.05 was considered statistically significant. Statistical analyses were performed using GraphPad Prism 9.0.

## 3. Results

### 3.1 Characterization of cultured EPCs

Cultured cells were characterized by DiI-ac-LDL uptake, FITC-UEA-1 binding, and EPC-related surface marker expression. As shown in Figure 1A, cells exhibited positive red fluorescence after DiI-ac-LDL incubation and positive green fluorescence after FITC-UEA-1 staining. The merged images confirmed the presence of double-positive cells, consistent with the functional phenotype of EPCs.

**Figure 1.**
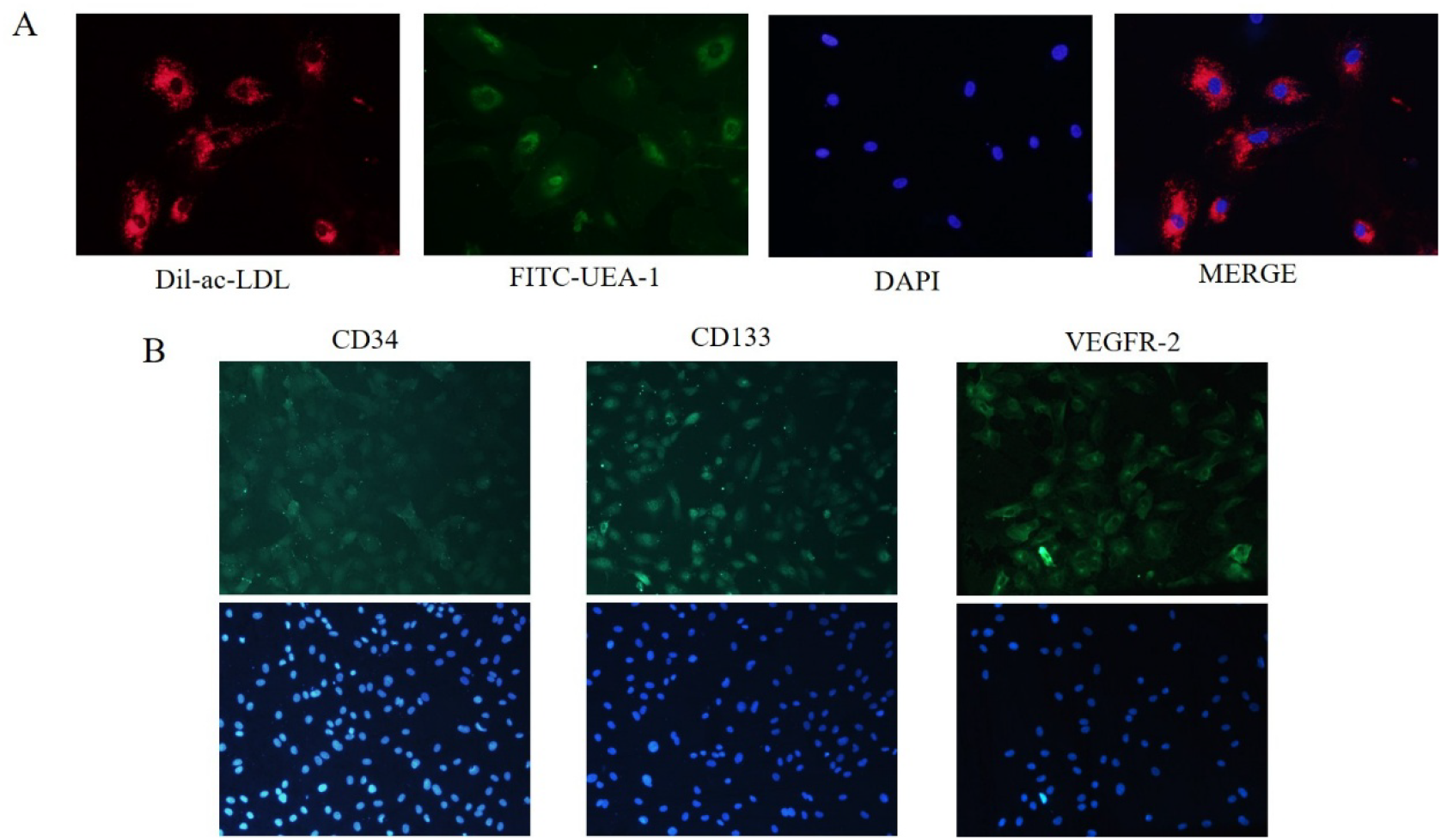
Characterization of cultured endothelial progenitor cells. (A) EPC identification by DiI-ac-LDL uptake and FITC-UEA-1 binding. Red fluorescence indicates DiI-ac-LDL uptake, green fluorescence indicates FITC-UEA-1 binding, and blue fluorescence indicates DAPI-stained nuclei. The merged image shows double-positive cells. (B) Immunofluorescence staining of EPC-related surface markers CD34, CD133, and VEGFR-2. Nuclei were counterstained with DAPI.

The EPC phenotype was further validated by immunofluorescence staining for CD34, CD133, and VEGFR-2. As shown in Figure 1B, the cultured cells were positive for all three EPC-related markers, with DAPI staining showing the corresponding nuclei. These results indicate that the cultured cells possessed typical EPC characteristics and were suitable for subsequent oxidative stress and functional assays.

### 3.2 H_2_O_2_ induces EPC apoptosis and impairs angiogenic capacity

To determine the effect of oxidative stress on EPC function, apoptosis and angiogenic capacity were evaluated by Annexin V/7-AAD flow cytometry and Matrigel tube formation assay, respectively. H_2_O_2_ treatment significantly increased EPC apoptosis in a concentration-dependent manner (Figure 2A). Compared with the control group, 200 μM H_2_O_2_ increased the apoptosis rate to 21.16 ± 3.13% (P < 0.01), while 400 μM and 500 μM H_2_O_2_ resulted in higher apoptosis rates of 31.86 ± 3.41% and 39.44 ± 3.61%, respectively (both P < 0.01 vs. control). Because 200 μM H_2_O_2_ produced reproducible injury without excessive cytotoxicity, this concentration was selected for subsequent intervention experiments.

**Figure 2.**
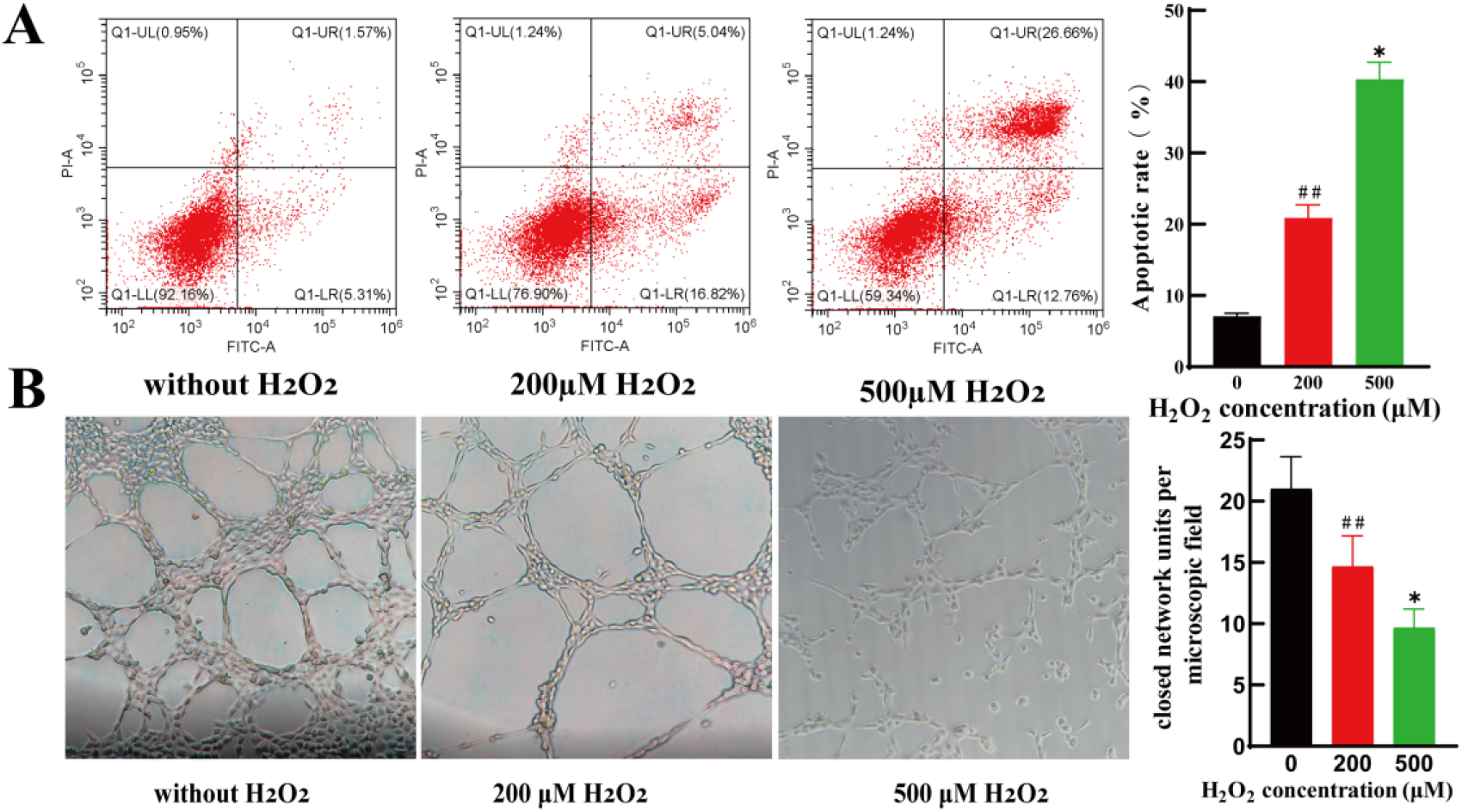
Effects of H_2_O_2_ on apoptosis and angiogenic capacity of EPCs. (A) EPC apoptosis after treatment with 0, 200, or 500 μM H_2_O_2_, analyzed by Annexin V-FITC/PI flow cytometry. Representative flow cytometry plots and quantitative analysis are shown. (B) Tube formation assay after H_2_O_2_ treatment. Representative images of capillary-like structures and quantitative analysis of closed network units are presented (magnification, ×100). Data are mean ± SD (n = 3). ##P < 0.01 vs. control group; *P < 0.05 vs. 200 μM H_2_O_2_ group.

The tube formation assay further showed that H_2_O_2_ impaired EPC angiogenic capacity (Figure 2B). Untreated EPCs formed well-organized and interconnected capillary-like networks. Exposure to 200 μM H_2_O_2_ reduced tube formation and disrupted network structure, whereas 500 μM H_2_O_2_ caused severe impairment, with sparse and discontinuous structures. These results demonstrate that H_2_O_2_ induces EPC dysfunction in a concentration-dependent manner.

### 3.3 H_2_O_2_ increases FoxO3a protein abundance and miR-34a expression in EPCs

The effects of oxidative stress on FoxO3a and miR-34a expression were examined by Western blot and qPCR. H_2_O_2_ reduced EPC viability in a concentration-dependent manner (Figure 3A). FoxO3a protein expression showed no obvious change at lower H_2_O_2_ concentrations (0-100 μM), but was significantly increased at 200 μM and further increased at 500 μM (Figure 3B). In contrast, FoxO3a mRNA levels remained relatively unchanged across the tested concentrations (Figure 3C), suggesting that oxidative stress may regulate FoxO3a primarily at the post-transcriptional or post-translational level.

**Figure 3.**
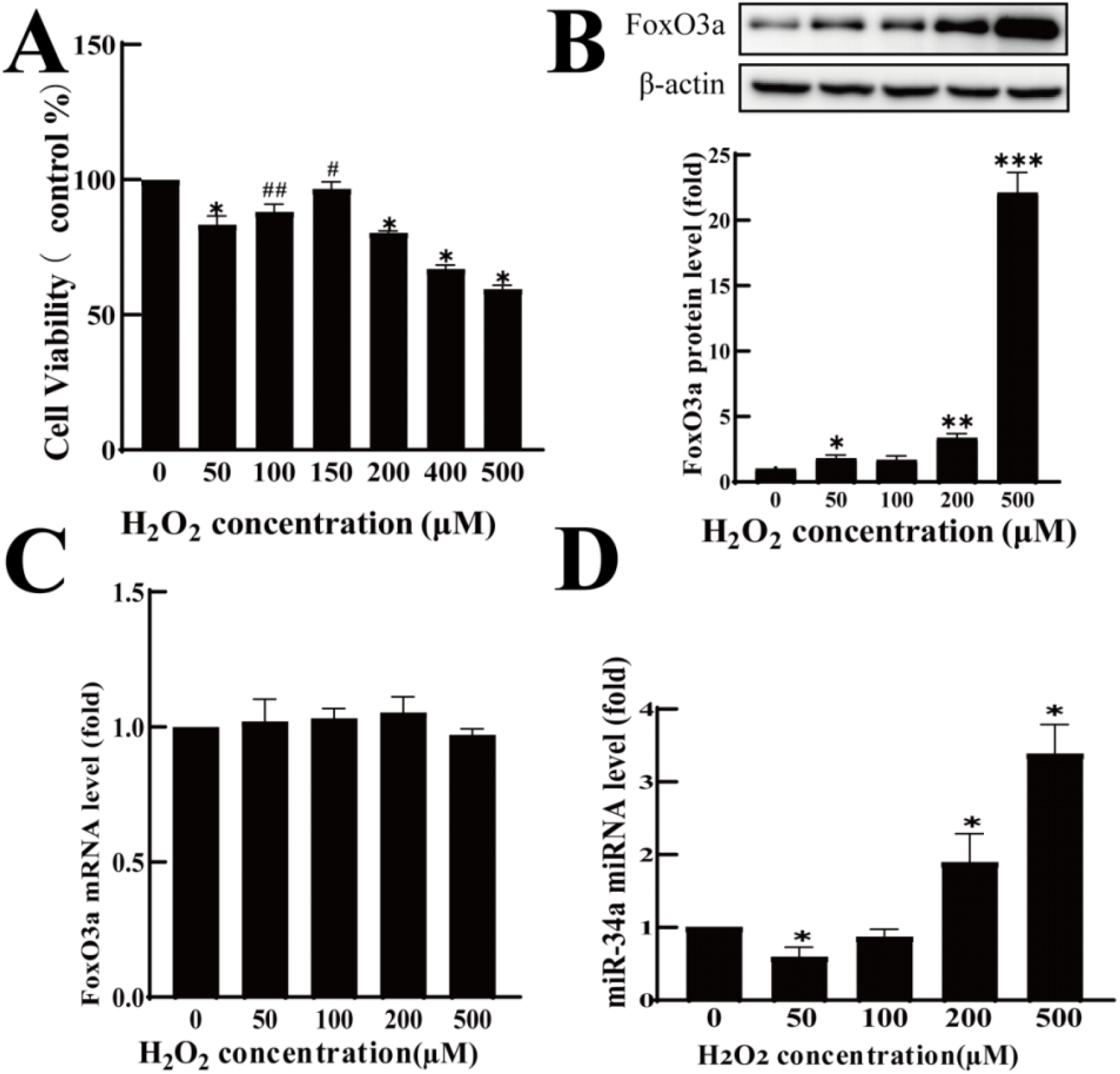
Effects of H_2_O_2_ on EPC viability and expression of FoxO3a and miR-34a. (A) EPC viability after exposure to increasing concentrations of H_2_O_2_, assessed by CCK-8 assay. (B) FoxO3a protein expression analyzed by Western blot and densitometric quantification. (C) FoxO3a mRNA expression measured by qPCR. (D) miR-34a expression measured by qPCR. Data are mean ± SD (n = 3).*P < 0.05, **P < 0.01, ***P < 0.001 vs. control group.

miR-34a expression did not differ markedly at lower H_2_O_2_ concentrations but increased significantly at concentrations of 200 μM and above (Figure 3D). These findings indicate that H_2_O_2_-induced oxidative stress is associated with increased FoxO3a protein abundance and elevated miR-34a expression in EPCs.

### 3.4 Optimization of adenoviral transduction efficiency in EPCs

To determine the optimal MOI for adenoviral transduction, EPCs were infected with adenoviruses at MOIs of 0, 5, 20, 40, 60, 80, 100, and 120. GFP fluorescence was detected in cells transduced with OE-NC, OE-FoxO3a, sh-NC, and sh-FoxO3a adenoviruses, confirming successful transduction (Figure 4). The proportion of GFP-positive cells and fluorescence intensity increased with increasing MOI.

**Figure 4.**
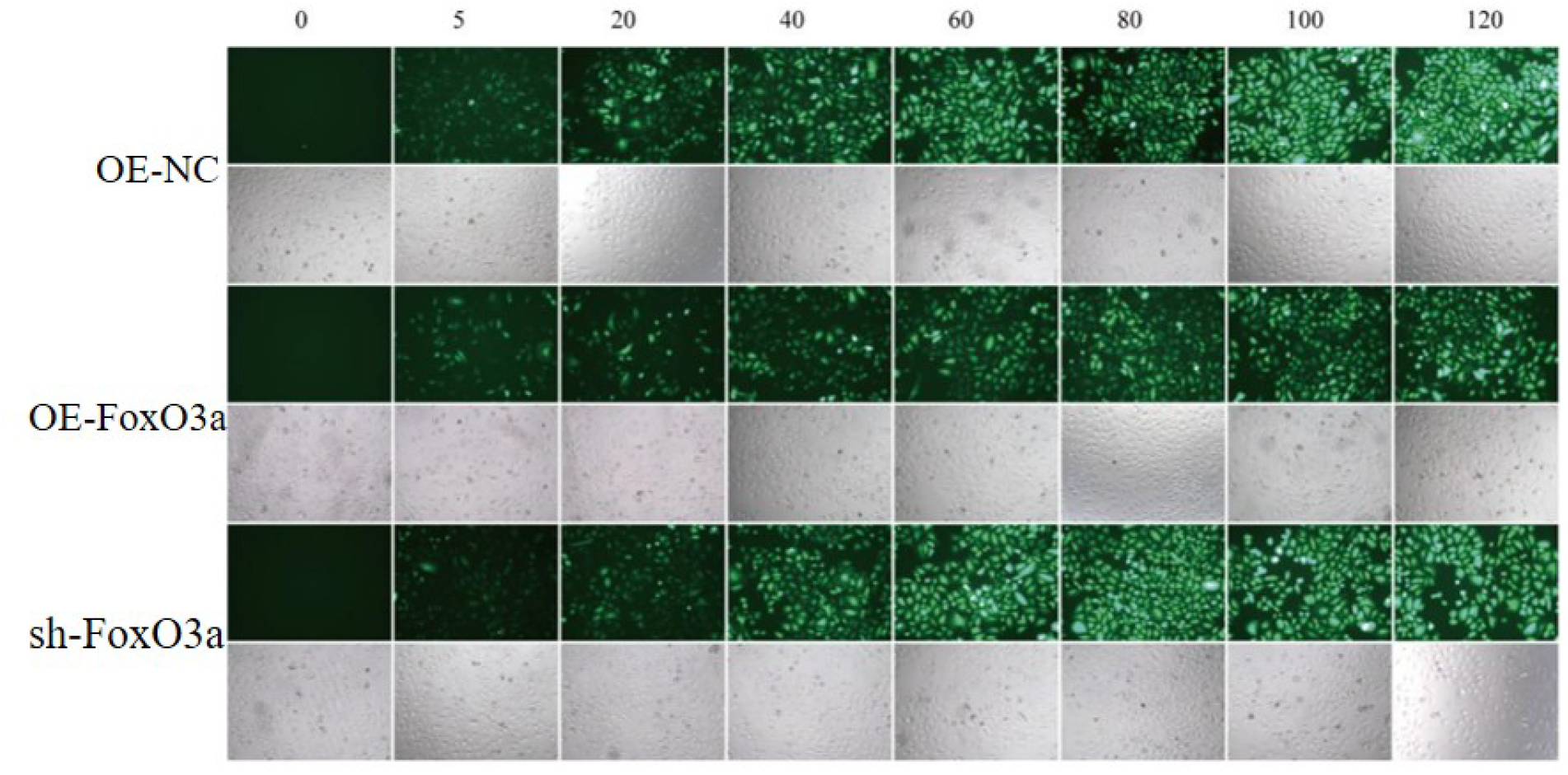
Optimization of adenoviral transduction efficiency in EPCs. Representative fluorescence images and corresponding bright-field images of EPCs transduced with control, FoxO3a overexpression, or FoxO3a knockdown adenoviruses at different MOIs (0-120), observed 48 h after transduction.

At an MOI of 120, however, obvious cytotoxicity was observed, including reduced cell density and altered cell morphology. Based on transduction efficiency and cell viability, an MOI of 80 was selected for sh-FoxO3a, while an MOI of 100 was used for OE-FoxO3a and control adenoviruses in subsequent experiments.

### 3.5 Validation of adenoviral and miRNA transfection efficiency

FoxO3a overexpression and knockdown efficiencies were validated by Western blot and qPCR. Compared with the corresponding control groups, OE-FoxO3a significantly increased FoxO3a protein and mRNA levels, whereas sh-FoxO3a markedly reduced FoxO3a expression (Figure 5A,B). These results confirmed successful adenoviral modulation of FoxO3a.

**Figure 5.**
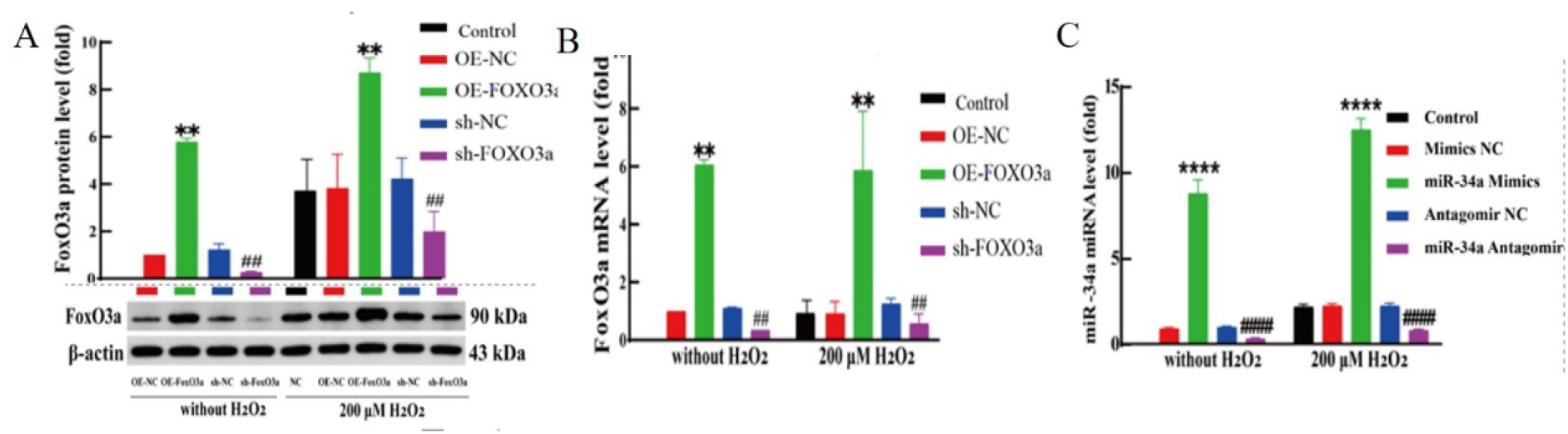
Validation of FoxO3a and miR-34a intervention efficiency in EPCs. (A) FoxO3a protein expression analyzed by Western blot and densitometric quantification in EPCs transduced with OE-FoxO3a, sh-FoxO3a, and corresponding controls. (B) FoxO3a mRNA expression measured by qPCR. (C) miR-34a expression measured by qPCR after transfection with miR-34a mimics, miR-34a antagomir, and corresponding negative controls. Data are mean ± SD (n = 3). **P < 0.01 vs. OE-NC group; ##P < 0.01 vs. sh-NC group; ****P < 0.0001 vs. mimics NC group; ####P < 0.0001 vs. antagomir NC group.

The efficiency of miR-34a mimic and inhibitor transfection was also confirmed by qPCR. miR-34a mimics significantly increased miR-34a expression compared with mimics NC, whereas miR-34a antagomir markedly reduced miR-34a expression compared with antagomir NC (Figure 5C). These results verified that the FoxO3a and miR-34a intervention systems were suitable for subsequent functional experiments.

### 3.6 FoxO3a and miR-34a regulate H_2_O_2_-induced reduction in EPC viability

The functional roles of FoxO3a and miR-34a in EPCs under oxidative stress were examined using the CCK-8 assay. H_2_O_2_ treatment significantly reduced EPC viability compared with the control condition (Figure 6A). FoxO3a overexpression further decreased EPC viability under H_2_O_2_ stimulation, whereas FoxO3a knockdown partially attenuated the H_2_O_2_-induced decline in viability.

**Figure 6.**
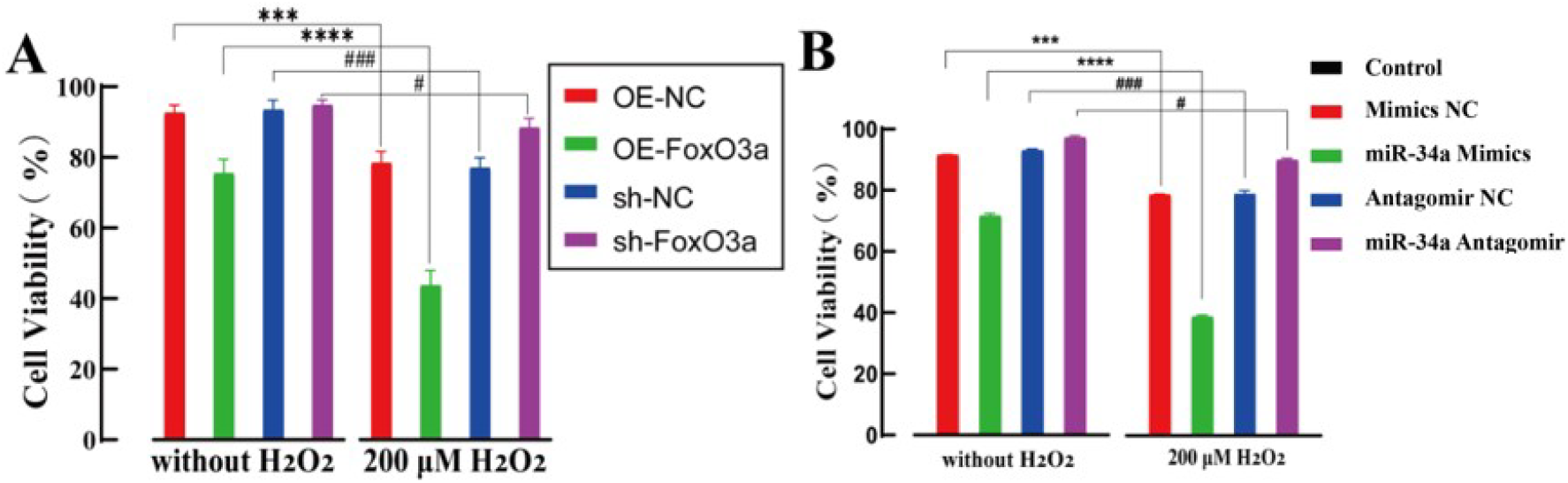
Effects of FoxO3a and miR-34a on EPC viability under oxidative stress. (A) Viability of EPCs transduced with OE-FoxO3a, sh-FoxO3a, and corresponding controls with or without 200 μM H_2_O_2_. (B) Viability of EPCs transfected with miR-34a mimics, miR-34a antagomir, and corresponding controls with or without 200 μM H_2_O_2_. Data are mean ± SD (n= 3). *P < 0.05, **P < 0.01, ***P < 0.001 vs. corresponding group without H_2_O_2_; #P < 0.05, ##P < 0.01, ###P < 0.001 vs. respective control group.

Modulation of miR-34a produced similar functional effects (Figure 6B). miR-34a mimics further reduced EPC viability under oxidative stress, while miR-34a antagomir significantly improved viability compared with the H_2_O_2_-treated negative control group. These findings indicate that both FoxO3a activation and miR-34a upregulation contribute to impaired EPC survival under oxidative stress.

### 3.7 FoxO3a and miR-34a regulate H_2_O_2_-induced impairment of EPC tube formation

The effects of FoxO3a and miR-34a on EPC angiogenic capacity were evaluated using an in vitro tube formation assay. EPCs transduced with OE-FoxO3a, OE-NC, sh-FoxO3a, or sh-NC were cultured on Matrigel with or without 200 μM H_2_O_2_. H_2_O_2_ treatment markedly reduced tube formation, as shown by the reduced number of closed network units (Figure 7A,B). FoxO3a overexpression further impaired tube formation, while FoxO3a knockdown significantly increased the number of closed network units under both basal and H_2_O_2_-treated conditions. These findings suggest that FoxO3a knockdown alleviates H_2_O_2_-induced angiogenic impairment in EPCs.

**Figure 7.**
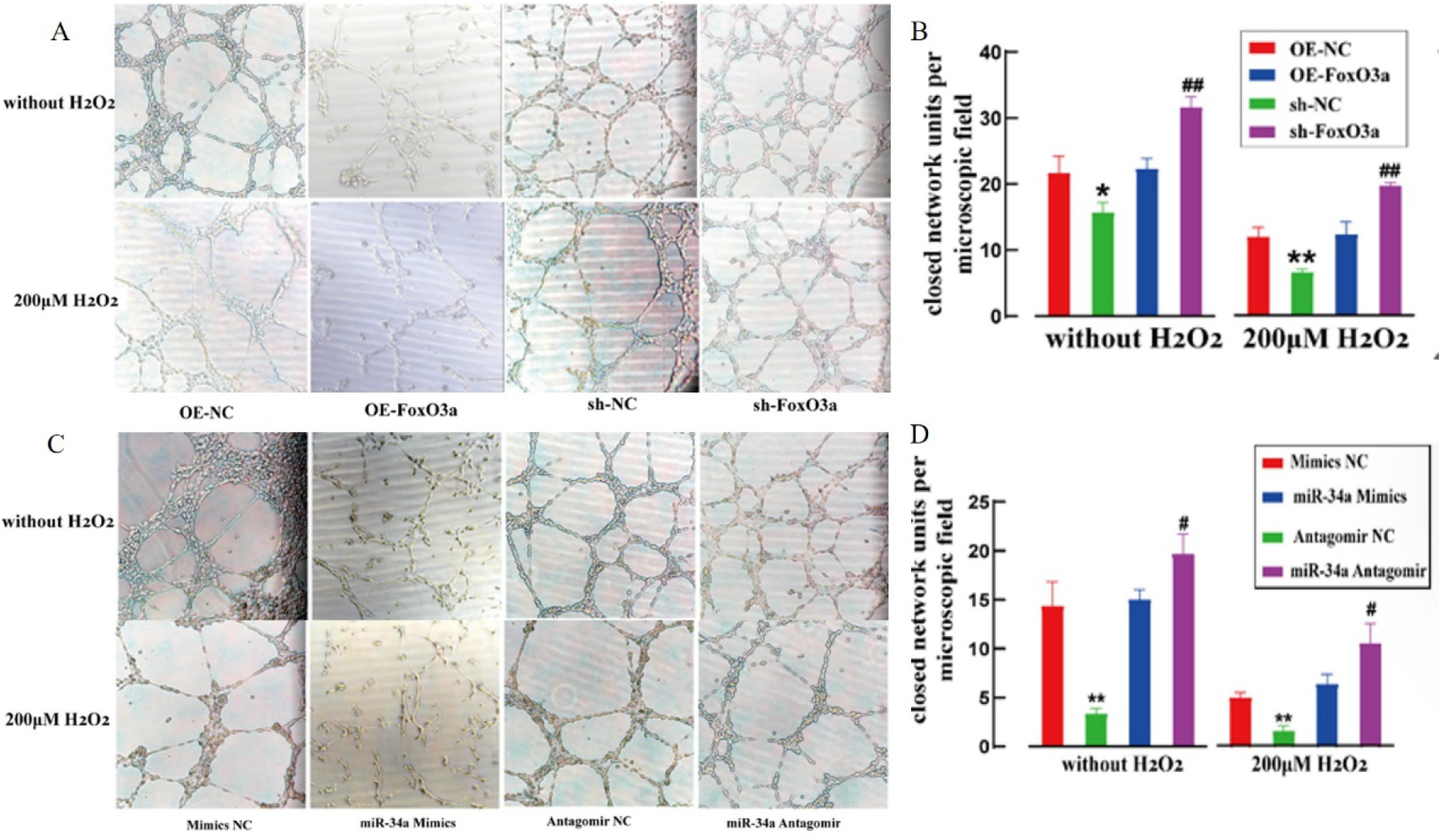
Effects of FoxO3a and miR-34a on H_2_O_2_-induced impairment of EPC tube formation in vitro. (A) Representative micrographs of EPCs transduced with OE-NC, OE-FoxO3a, sh-NC, or sh-FoxO3a and cultured on Matrigel with or without 200 μM H_2_O_2_ for 24h. (B) Quantification of closed network units in the FoxO3a intervention groups. (C) Representative micrographs of EPCs transfected with mimics NC, miR-34a mimics, antagomir NC, or miR-34a antagomir and cultured on Matrigel with or without 200 μM H_2_O_2_ for 24 h. (D) Quantification of closed network units in the miR-34a intervention groups. Magnification, ×100. Data are mean ± SD (n = 3). *P < 0.05 and **P < 0.01 vs. corresponding NC group; #P < 0.05 and ##P < 0.01 vs. corresponding NC group.

The effect of miR-34a on EPC angiogenic capacity was then assessed. miR-34a mimics significantly reduced closed network formation, whereas miR-34a antagomir improved tube formation, especially under H_2_O_2_ stimulation (Figure 7C,D). These results indicate that miR-34a aggravates oxidative stress-induced angiogenic dysfunction in EPCs, while miR-34a inhibition exerts a protective effect.

### 3.8 hsa-miR-34a-3p directly targets the FoxO3a 3′UTR

To verify whether miR-34a can directly regulate FoxO3a at the post-transcriptional level, dual-luciferase reporter assays were performed using pmirGLO constructs containing either the wild-type FoxO3a 3′UTR or a mutant sequence with disrupted hsa-miR-34a-3p seed-matching nucleotides (Figure 8A).

**Figure 8.**
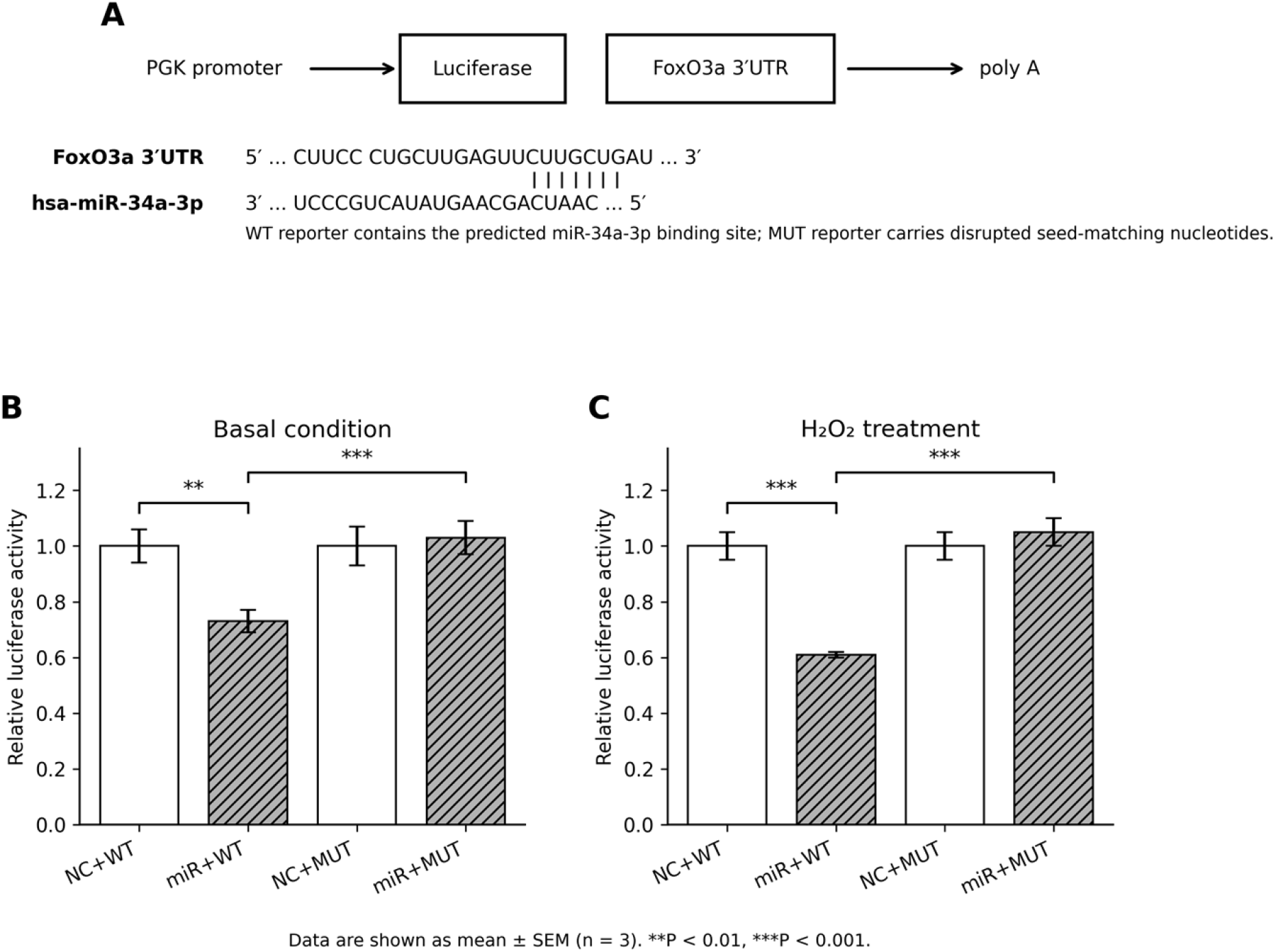
hsa-miR-34a-3p directly binds the FoxO3a 3′UTR and suppresses reporter activity. (A) Schematic of the predicted hsa-miR-34a-3p binding site in the FoxO3a 3′UTR and corresponding mutant reporter design. (B) Relative luciferase activity in 293T cells co-transfected with WT or MUT reporter plasmids and hsa-miR-34a-3p mimic or negative control under basal conditions. (C) Relative luciferase activity after H_2_O2 treatment. Firefly luciferase activity was normalized to Renilla luciferase activity. Data are mean ± SEM (n = 3). **P < 0.01, ***P < 0.001.

Under basal conditions, hsa-miR-34a-3p significantly decreased normalized luciferase activity of the FoxO3a 3′UTR-WT reporter to 0.73 ± 0.04 relative to the negative control group (P = 0.0056; Figure 8B). Mutation of the predicted binding site abolished this suppression, and luciferase activity remained comparable to the mutant negative control group (1.03 ± 0.04 vs. 1.00 ± 0.04; P > 0.05).

After H_2_O_2_ treatment, hsa-miR-34a-3p further reduced the activity of the FoxO3a 3′UTR-WT reporter to 0.61 ± 0.01 relative to the corresponding negative control (P = 0.0002; Figure 8C). Again, mutation of the binding site eliminated the inhibitory effect of hsa-miR-34a-3p. These results confirm that hsa-miR-34a-3p directly targets the FoxO3a 3′UTR and suggest that miR-34a-3p-mediated post-transcriptional feedback may participate in FoxO3a regulation during oxidative stress.

## 4. Discussion

Oxidative stress is a major contributor to endothelial dysfunction and impaired vascular repair. In this study, H_2_O_2_ exposure significantly compromised EPC function, as shown by reduced cell viability, increased apoptosis, and impaired tube formation. These changes were accompanied by increased FoxO3a protein abundance and elevated miR-34a expression. Functional intervention experiments further showed that FoxO3a overexpression aggravated EPC injury, whereas FoxO3a knockdown partially alleviated H_2_O_2_-induced dysfunction. Similar trends were observed after miR-34a modulation: miR-34a mimics further impaired EPC function, whereas miR-34a antagomir exerted a protective effect. Collectively, these findings indicate that FoxO3a and miR-34a are involved in oxidative stress-induced EPC dysfunction.

The concentration-dependent injury induced by H_2_O_2_ is consistent with the established role of ROS in endothelial damage. Excessive ROS accumulation reduces nitric oxide bioavailability, disrupts mitochondrial homeostasis, promotes inflammatory signaling, and triggers apoptosis [11,12]. In the present study, 200 μM H_2_O_2_ caused significant but not overwhelming injury, allowing the effects of genetic and miRNA interventions to be evaluated. At higher concentrations, severe cytotoxicity may obscure regulatory effects by inducing extensive cell death. Therefore, the use of 200 μM H_2_O_2_ provided a stable injury model for assessing the contributions of FoxO3a and miR-34a to EPC dysfunction.

FoxO3a showed a detrimental role in this H_2_O_2_-induced injury model. Overexpression of FoxO3a reduced EPC viability and tube formation, whereas FoxO3a knockdown improved these functional parameters. FoxO3a has complex and context-dependent effects in stress biology. Although it may support antioxidant adaptation under mild stress, sustained activation can promote apoptosis and inhibit regenerative capacity [5,14]. Previous work has linked H_2_O_2_-induced EPC viability impairment to FoxO3a-dependent mechanisms [4], and our results extend this evidence by showing that FoxO3a also contributes to angiogenic impairment under oxidative stress.

A notable finding was that H_2_O_2_ increased FoxO3a protein abundance without a corresponding increase in FoxO3a mRNA. This discrepancy suggests that FoxO3a regulation under oxidative stress occurs predominantly at the post-transcriptional or post-translational level. FoxO3a activity is controlled by phosphorylation, acetylation, ubiquitination, protein stability, subcellular localization, and interactions with other signaling molecules. Oxidative stress may enhance FoxO3a protein stability, reduce degradation, or promote nuclear retention even in the absence of increased mRNA expression. Future studies should examine phosphorylated FoxO3a, nuclear-cytoplasmic distribution, and ubiquitination status to better define the activation mode of FoxO3a in EPCs.

miR-34a also acted as a negative regulator of EPC function in this study. miR-34a mimics reduced EPC viability and tube formation, whereas miR-34a antagomir partially restored these functions under oxidative stress. These results are consistent with reports linking miR-34a to endothelial senescence, apoptosis, impaired angiogenesis, and age-related vascular dysfunction [7,15]. Because EPC function is regulated by multiple non-coding RNAs, the partial protection produced by miR-34a inhibition likely reflects the contribution of a broader regulatory network that may include miR-221-3p, miR-29c-3p, miR-126a, and other EPC-associated microRNAs [16-18].

The dual-luciferase reporter assay clarified an important mechanistic point. The data demonstrate that hsa-miR-34a-3p directly targets the FoxO3a 3′UTR, because hsa-miR-34a-3p significantly suppressed the activity of the WT reporter, whereas mutation of the predicted binding site abolished this inhibitory effect. This result should not be interpreted as evidence that FoxO3a activates the miR-34a promoter. Instead, it supports a model in which miR-34a-3p can post-transcriptionally regulate FoxO3a through direct 3′UTR binding. The simultaneous increase in FoxO3a protein and miR-34a expression after H_2_O_2_ exposure suggests a more complex regulatory circuit: oxidative stress may increase FoxO3a protein stability or activity, while miR-34a-3p-mediated 3′UTR targeting may provide a negative-feedback mechanism that restrains FoxO3a expression.

SIRT1 remains a biologically relevant component of this network. SIRT1 protects endothelial cells and EPCs by regulating oxidative stress, mitochondrial homeostasis, inflammation, apoptosis, and angiogenesis [6,9,19,20]. miR-34a has been reported to repress SIRT1 in vascular contexts [7,15], and SIRT1 can modulate FoxO3a stability and activity [6]. Thus, the FoxO3a/miR-34a relationship identified in this study may interact with SIRT1-associated protective pathways. However, direct confirmation of a causal miR-34a/SIRT1/FoxO3a axis in EPCs will require additional rescue experiments, such as SIRT1 overexpression or pharmacological activation combined with miR-34a and FoxO3a interventions.

This study has several limitations. First, the experiments were performed mainly in an in vitro H_2_O_2_-induced oxidative stress model. Although this model is useful for mechanistic analysis, it does not fully reproduce the complexity of vascular injury in vivo. Second, the dual-luciferase reporter assay was performed in 293T cells, which provide high transfection efficiency but are not EPCs; therefore, validation in EPCs or endothelial-lineage cells would strengthen the conclusion. Third, although functional data support the involvement of FoxO3a and miR-34a in EPC dysfunction, additional mechanistic rescue experiments are required to define the precise hierarchy among FoxO3a, miR-34a, SIRT1, and downstream apoptosis-related pathways. Finally, animal models of vascular injury, diabetes, or atherosclerosis are needed to evaluate whether targeting the FoxO3a/miR-34a regulatory network improves EPC-mediated vascular repair in vivo.

In summary, H_2_O_2_-induced oxidative stress impairs EPC survival and angiogenic capacity and is accompanied by increased FoxO3a protein abundance and miR-34a expression. FoxO3a overexpression and miR-34a mimic transfection aggravate EPC dysfunction, whereas FoxO3a knockdown and miR-34a inhibition exert protective effects. The dual-luciferase reporter assay confirms that hsa-miR-34a-3p directly targets the FoxO3a 3′UTR, indicating that miR-34a-3p-mediated post-transcriptional feedback is part of the FoxO3a-related stress-response network. These findings provide a mechanistic basis for further investigation of FoxO3a and miR-34a as potential targets for improving EPC-mediated vascular repair under oxidative stress.

## 5. Conclusion

FoxO3a and miR-34a contribute to H_2_O_2_-induced EPC dysfunction. FoxO3a promotes oxidative stress-related reductions in EPC viability and angiogenic capacity, while miR-34a aggravates these impairments and directly targets the FoxO3a 3′UTR. These findings support a regulatory network in which FoxO3a-associated stress responses and miR-34a-3p-mediated post-transcriptional feedback jointly influence EPC injury under oxidative stress.

## Ethics statement

Human umbilical cord blood samples were obtained from donors who provided informed consent. The study was approved by the institutional ethics committee.

